# Induction of SARS-CoV-2 N-specific CD8^+^ T cell immunity in lungs by engineered extracellular vesicles associates with strongly impaired viral replication

**DOI:** 10.1101/2023.01.19.524762

**Authors:** Francesco Manfredi, Chiara Chiozzini, Flavia Ferrantelli, Patrizia Leone, Katherina Pugliese, Massimo Spada, Antonio Di Virgilio, Andrea Giovannelli, Mauro Valeri, Andrea Cara, Zuleika Michelini, Mauro Andreotti, Maurizio Federico

## Abstract

Induction of effective immunity in lungs should be a pre-requisite for any vaccine designed to control the severe pathogenic effects generated by respiratory infectious agents. In the case of Severe Acute Respiratory Syndrome Coronavirus (SARS-CoV)-2 infection, vaccination is expected to associate with significant inhibition of viral replication in lungs. We recently provided evidence that the generation of endogenous extracellular vesicles (EVs) engineered for the incorporation of SARS-CoV-2 Nucleocapsid (N) protein can protect K18-hACE2 transgenic mice from the lethal intranasal infection with the ancestral Wuhan isolate. Actually, it was widely demonstrated that these transgenic mice succumb to SARS-CoV-2 intranasal infection mainly as a consequence of the viral invasiveness of central nervous system, a pathogenetic mechanism almost absent in humans. On the other hand, K18-hACE2 transgenic mice support viral replication in lungs, an event strictly mirroring the major pathogenic signature linked to the severe disease in humans. However, nothing is known about the ability of N-specific CD8^+^ T cell immunity induced by engineered EVs in controlling viral replication in lungs. To fill the gap, we investigated the immunity generated in lungs by N-engineered EVs in terms of induction of N-specific effectors and resident memory CD8^+^ T lymphocytes before and after virus challenge carried out three weeks and three months after boosting. At the same time points, viral replication extents in lungs were evaluated. We found that three weeks after second immunization, virus replication was reduced in mice best responding to vaccination by more than 3-logs compared to control group. The impaired viral replication matched with a reduced induction of Spike-specific CD8^+^ T lymphocytes. The antiviral effect appeared similarly strong when the viral challenge was carried out 3 months after boosting. This inhibitory effect associated with the persistence of a N-specific CD8^+^ T-resident memory lymphocytes in lungs of N-immunized mice. In view of the quite conserved sequence of the N protein among SARS-CoV-2 variants, these results support the idea that a vaccine strategy focused on the induction of anti-N CD8^+^ T cell immunity in lungs has the potential to control the replication of emerging variants.

## Introduction

Several vaccines were developed and distributed within an unprecedented short time in response to the spread of Severe Acute Respiratory Syndrome Coronavirus (SARS-CoV)-2. All vaccine preparations were conceived to elicit anti-Spike protein immune responses, and their effectiveness relies on the generation of neutralizing antibodies. Anti-SARS-CoV-2 mRNA-based vaccines lead to the production of extraordinarily high levels of anti-Spike antibodies in serum [1,2]. Regrettably, however, the presence of Spike-binding antibodies in humans at the viral port of entry, i.e., the mucosa of the upper respiratory tract, does not result in a significant virus neutralization [3-7]. In addition, anti-Spike immunity in lungs of vaccinees was found at levels of the detection threshold [8].

The almost absence of Spike-specific cell immunity in lungs of vaccinees is not so surprising since it is known that the development of cell immunity in pulmonary tissues is largely not influenced by events occurring in both peripheral circulation and distal lymphoid organs. Lymphocytes in lungs are maintained independently of the pool of circulating lymphocytes, and their continuous loss through intraepithelial migration towards the airways is constantly replenished by homeostatic proliferation [9]. In this scenario, the identification of new anti-SARS-CoV-2 vaccines eliciting adequate antiviral immunity in lungs would be of outmost relevance.

All cell types constitutively release nanovesicles (collectively referred to as extracellular vesicles, EVs), which are key players in intercellular communication [10]. We developed a CD8^+^ T-cell based vaccine platform based on intramuscular (i.m.) injection of a DNA vector coding for antigens of interest fused at the C-terminus of a biologically inactive Human Immunodeficiency Virus (HIV)-Type 1 Nef protein (Nef^mut^) having an unusually high efficiency of incorporation into EVs. This unique feature is maintained even when foreign polypeptides are fused to its C-terminus [11, 12]. Both N-terminal myristoylation and palmitoylation fasten Nef^mut^ to the luminal membrane leaflets, hence resulting critical for its abundant uploading in EVs. Upon i.m. injection of DNA vectors expressing Nef^mut^-derivatives, nanovesicles containing antigens fused with Nef^mut^ are released by muscle cells, and can freely circulate into the body and be internalized by antigen-presenting cells (APCs). EV-associated antigens are then cross-presented to prime antigen-specific CD8^+^ T lymphocytes [13].

We already tested the efficacy of the Nucleocapsid (N)-specific CD8^+^ T cell immunity elicited by EVs engineered *in vivo* for the incorporation of SARS-CoV-2 protein in survival assays carried out on K18-hACE2 transgenic mice [14]. These mice replicate SARS-CoV-2 by virtue of the expression of the human ACE2 receptor expressed under the control of a human cytokeratin-18 promoter which allows viral entry in epithelial cells [15]. However, it was reproducibly shown that the heavy pathogenic effects observed in infected mice essentially rely on viral invasion of central nervous system, while the infection in lung tissues contributes to the overall health decay and weight loss only marginally [16-19]. Nevertheless, SARS-CoV-2 replicates in lungs of K18-hACE2 mice as efficiently as in those of infected humans [16]. For these reasons, we were interested in evaluating the efficacy of the EV-induced N-specific immunity in counteracting the viral replication in lungs. To this end, both CD8^+^ T cell immunity and viral loads in lungs were measured in mice challenged 3 weeks and 3 months after immunization. The strong viral inhibition we observed in lungs of N-immunized mice paves the way towards the design of second-generation anti-SARS-CoV-2 vaccines which, considering the extremely low variability of N protein [14], would be effective against emerging variants.

## Materials & Methods

### DNA constructs

Open-reading frames coding for either Nef^mut^ alone or fused with SARS-CoV-2 N protein were cloned into pVAX1 plasmid (Thermo Fisher, Waltham, MA) as previously described [20]. In the Nef^mut^/N construct, the entire open reading frame for N protein (422 aa), except the M_1_ amino acid, is part of the fusion product. A GPGP linker was inserted between Nef^mut^ and the N sequences. Stop codon was preceded by sequences coding for a DYKDDDK epitope tag (flag-tag). SARS-CoV-2 sequences were optimized for expression in human cells through GenSmart™ Codon Optimization software from Genescript (Piscataway, NJ). All vectors were synthesized by OfficinaeBio (Venice, Italy).

### Animals and authorizations

Six-week-old C57 Bl/6 K18-hACE2 transgenic female mice were purchased from Charles River (Calco, Italy) and hosted at the Central and BSL3 Animal Facilities of the Istituto Superiore di Sanità, as approved by the Italian Ministry of Health, authorizations 565/2020 and 591/2021, and released on June 3rd 2020 and July 30th 2021, respectively. Before the first procedure, Datamars (Lugano, Switzerland) microchips were inserted subcutaneously on the dorsal midline between the shoulder blades.

### Mice immunization

Isoflurane-anesthetized mice were inoculated i.m. with 10 μg of DNA in 30 μL of sterile, 0.9% saline solution. DNA injection was immediately followed by electroporation at the site of inoculation with an Agilpulse BTX (Holliston, MA) device, using a 4-needle electrode array (4 mm gap, 5 mm needle length) and applying the following parameters: 1 pulse of 450 V for 50 μs; 0.2 ms interval; 1 pulse of 450 V for 50 μs; 50 ms interval; 8 pulses of 110 V for 10 ms with 20 ms intervals. Mice were immunized into both quadriceps, twice, 2 weeks apart. Mice were sacrificed by cervical dislocation.

### IFN-γ EliSpot analysis

Either 2.5 (for splenocytes) or 1 (for peripheral blood mononuclear cells, PBMCs)×10^5^ live cells were seeded in triplicate in microwells of 96-multiwell plates (Millipore, Burlington, MA) previously coated with the anti-mouse IFN-AN18 mAb (Mabtech, Nacka Strand, Sweden) in RPMI 1640, 10% heat-inactivated fetal calf serum (FCS, Gibco, Thermo Fisher), and 50 μM 2-mercaptoethanol. Cell cultures were carried out for 16 h in the presence of 5 μg/mL of CD8-specific H2-b-binding SARS-CoV-2 Spike: 539–546 VNFNFNGL [21], or N peptides: 219–228 ALALLLLDRL [22], or CD8-specific H2-b-binding HIV-1 Nef peptide: 48-56 TAATNADCA [23]. As negative controls, 5 μg/mL of unrelated H2-b binding peptides were used. Peptide preparations were obtained from BEI resources. To check for cell responsiveness, 10 ng/mL phorbol 12-myristate 13-acetate (PMA, Sigma, St. Louis, MO) plus 500 ng/mL of ionomycin (Sigma) were added to the cultures. After 16 h, the cells were discarded, and the plate was incubated for 2 h at room temperature with R4-6A2 biotinylated anti-IFN-γ antibody (Mabtech) at the concentration of 100 μg/mL. Wells were then washed and treated for 1 h at room temperature with 1:1,000 diluted streptavidin-ALP from Mabtech. Afterwards, 100 μL/well of SigmaFast BCIP/NBT were added to the wells to develop spots. Spot-forming cells were finally analyzed and counted using an AELVIS EliSpot reader (Hannover, Germany).

### Cell isolation from spleens, blood, and lungs

PBMCs were recovered from EDTA-blood samples obtained through retro orbital puncture under topical anesthesia. Erythrocytes were removed by treatment with ACK lysing buffer (Gibco) according to the manufacturer’s instructions.

To isolate splenocytes, spleens were explanted, placed into tubes containing 1 mL of RPMI 1640 and 50 µM 2-mercaptoethanol, then transferred into 60 mm Petri dishes with 2 mL of the same medium. Splenocytes were obtained by notching the spleen sac and pushing the cells out with the plunger seal of a 1 mL syringe. After addition of 2 mL of medium, cells were transferred into a 15 mL conical tube, and the Petri plate washed with 4 mL of medium to maximize cell recovery. Afterwards, cells were collected by centrifugation, resuspended in RPMI complete medium containing 50 µM 2-mercaptoethanol and 10% FCS, and counted.

For lung cell isolation, circulating blood cells were fluorescently labeled with 2 μg (10 µl of the commercial stock) of an anti-CD45 antibody (Tonbo-Bioscience, S. Diego, CA anti-mouse CD45.2-APC) diluted in 200 µl of 1×PBS inoculated in the tail vein exactly 3 minutes before cervical dislocation. For cell recovery, lungs were excised, washed with 1×PBS, cut into small pieces, and then digested for 30 minutes under gentle agitation at 37°C with 7 mL of 4 mg/mL of type III collagenase (Worthington Biochemical, Lakewood, NJ) and 0.05 mg/mL of DNase I (Sigma) in 1×PBS. After digestion, an equal volume of medium was added and cells were passed through a 70 µm cell strainer, washed, and resuspended in 1×PBS/ACK 1:1 for red blood cells lysis. Finally, the isolated cells were resuspended in complete culture medium and counted.

### Intracellular cytokine staining (ICS) and flow cytometry analysis

Cells collected from blood, spleens, and lungs were cultured at 1×10^7^/mL in RPMI medium, 10% FCS, 50 µM 2-mercaptoethanol (Sigma), 1 µg/mL brefeldin A (BD Biosciences, Franklin Lakes, NJ), and in the presence of 5 μg/mL of either Spike, N, or unrelated H2-b CD8^+^ T-specific peptides. Positive controls were conducted by adding 10 ng/mL PMA (Sigma) plus 1 µg/mL ionomycin (Sigma). After 16 h, cells were stained with 1 µL of LIVE/DEAD Fixable FVD-eFluor506 Dead Cell reagent (Invitrogen Thermo Fisher) in 1 mL of 1×PBS for 30 min at 4 °C, and excess dye removed by 2 washes with 500 µL of 1×PBS. Non-specific staining was minimized by pre-incubating cells with 0.5 µg of Fc blocking mAbs (i.e., anti-CD16/CD32 antibodies, Invitrogen/eBioscience Thermo Fisher) in 100 µL of 1×PBS with 2% FCS for 15 min at 4 °C. Staining for cell surface markers was performed upon incubation for 1 h at 4 °C with 2 µL of the following anti-mouse Abs: FITC-conjugated anti-CD3, APC-Cy7-conjugated anti-CD8a, PerCP-conjugated anti-CD4, and BUV395-conjugated anti-CD44 (BD Biosciences). CD8^+^ T-resident memory (Trm) cells were identified by staining with BUV750-conjugated anti-CD49a, PECF594-conjugated anti-CD69, and BUV563-conjugated anti-CD103 (BD Biosciences). For intracellular cytokine staining (ICS), cells were fixed and permeabilized using the Cytofix/Cytoperm kit (BD Biosciences), according to the manufacturer’s recommendations. Thereafter, cells were labeled for 1 h at 4 °C with 2 µL of the following Abs: PE-Cy7-conjugated anti-IFN-γ, PE-conjugated anti-IL-2 (Invitrogen/eBioscience Thermo Fisher), and BV421 rat anti-TNF-α (BD Biosciences) in a total of 100 µL of 1× Perm/Wash Buffer (BD Biosciences).

After two washes, cells were fixed in 200 µL of 1× PBS/formaldehyde (2% v/v). Samples were then analyzed by a CyotFLEX LX (Beckman Coulter, Brea, CA, USA) flow cytometer and analyzed using Kaluza software (Beckman Coulter). Gating strategy was as follows (Supplementary figs.1 and 2): live cells as assessed by LIVE/DEAD dye vs. FSC-A, singlet cells from FSC-A vs. FSC-H (singlet 1) and SSC-A vs. SSC-W (singlet 2), CD3^+^ cells from CD3-FITC vs. SSC-A, CD8^+^, or CD4^+^ cells from CD8-APC-Cy7 vs. CD4-PerCP. CD3+/CD8^+^ cell population was gated against CD44^+^ cells, and, to detect polyfunctional CD8^+^ T lymphocytes, the population of cells positive for both CD8 and CD44 was analyzed for APC-Cy7, PE, and BV421 to detect simultaneous changes in IFN-γ, IL-2, and TNF-α production, respectively. To detect CD8^+^ T resident memory (Trm), the population of cells positive for both CD8 and CD44 was analyzed for the co-expression of CD49a, CD69, and CD103. Boolean gates were created to measure co-expression patterns.

### Virus production

VERO-E6 cells were grown in DMEM (Gibco, Thermo Fisher) supplemented with 2% FCS, 100 units/mL penicillin, 100 μg/mL streptomycin, 2 mM L-glutamine, 1 mM sodium pyruvate, and non-essential amino acids (Gibco). The ancestral viral isolate SARS-CoV-2/Italy INMI1#52284 (SARS-Related Coronavirus 2, isolate Italy-INMI1, NR-52284, deposited by Dr. Maria R. Capobianchi for distribution through BEI Resources, NIAID, and NIH) was propagated by inoculation of 70% confluent VERO-E6 cells [24]. Infected cell culture supernatant was harvested at 72 h post infection, clarified, aliquoted, and stored at -80 °C.

### Mouse infection

Before experimental infection, mice were anesthetized with a combination of ketamine (50 mg/kg of body weight) and medetomidine (1 mg/kg of body weight) administered intraperitoneally. A volume of 30 μL of each dilution was administered intranasally (i.n.), at 15 μL per nostril, dropwise. After virus challenge, intraperitoneal injection of atipamezole (1 mg/kg of body weight) was used as a reversal agent. For *in vivo* titration of the SARS-CoV-2/Italy INMI1#52284 isolate, age-matched K18-hACE2 mice were randomized by body weight into groups of 4, and challenged with 5-fold serial dilutions in 1×PBS of the virus preparation containing 2.2×10^5^, 4.4 10^4^, 8.8×10^3^, or 1.8×10^3^ TCID_50_ measured as previously reported [14]. As a negative control, 4 mice were sedated and an equal volume of 1×PBS was administered. Animals were assessed daily for clinical signs of infection. Virus challenge in immunized mice was performed using a virus dose of 4.4 lethal doses at 50% (LD_50_) resulting in a 99.99% predicted probability of mortality in untreated mice.

### Extraction and purification of lung RNA

Lungs were stored frozen after excision. After thawing, equal amounts of tissues were minced and incubated for 10 min in 1 mL of TRIzol™ Reagent (Thermo Fisher Scientific). Minced tissue was then passed through a QIAshredder homogenizer (Qiagen, Germantown, MD), and the flow-through was used for chloroform extraction according to the TRIzol™ protocol, using 0.2 mL of chloroform. Recovered total RNA was stored in water at -80 °C.

### RT-qPCR

RT-qPCR for SARS-CoV-2 Envelope (E) and Nucleocapsid (N) genes was performed using a One-Step Taqman-based strategy. Mouse β-actin amplification was included as loading control. All probes and primers (Table 1) were purchased from Integrated DNA Technologies (IDT). Each 20 µL reaction mixture contained 12 μL of qPCRBIO Probe 1-Step Virus Detect Lo ROX master mix (PCR Biosystems Ltd.London, UK), 3 µL of primers/probes mix, and of 1 µg of ezDNAse (Thermo Fisher)-treated RNA in a total of 5 µL. All samples were tested in duplicate, and samples with nuclease-free water alone were included as negative controls. Serial 10-fold dilutions of E gene plasmid (10006896, 2019-nCoV_E Positive Control from Charité/Berlin, IDT, Leuven, Belgium) and N1/N2 plasmid (10006625, 2019-nCoV_N_Positive Control from CDC, IDT, Belgium) were used to generate standard curves ranging from 1 to 100,000 copies. Median standard curve slope ranged from 3.25 to 3.43 with R^2^>0.998.

**Table 1.**
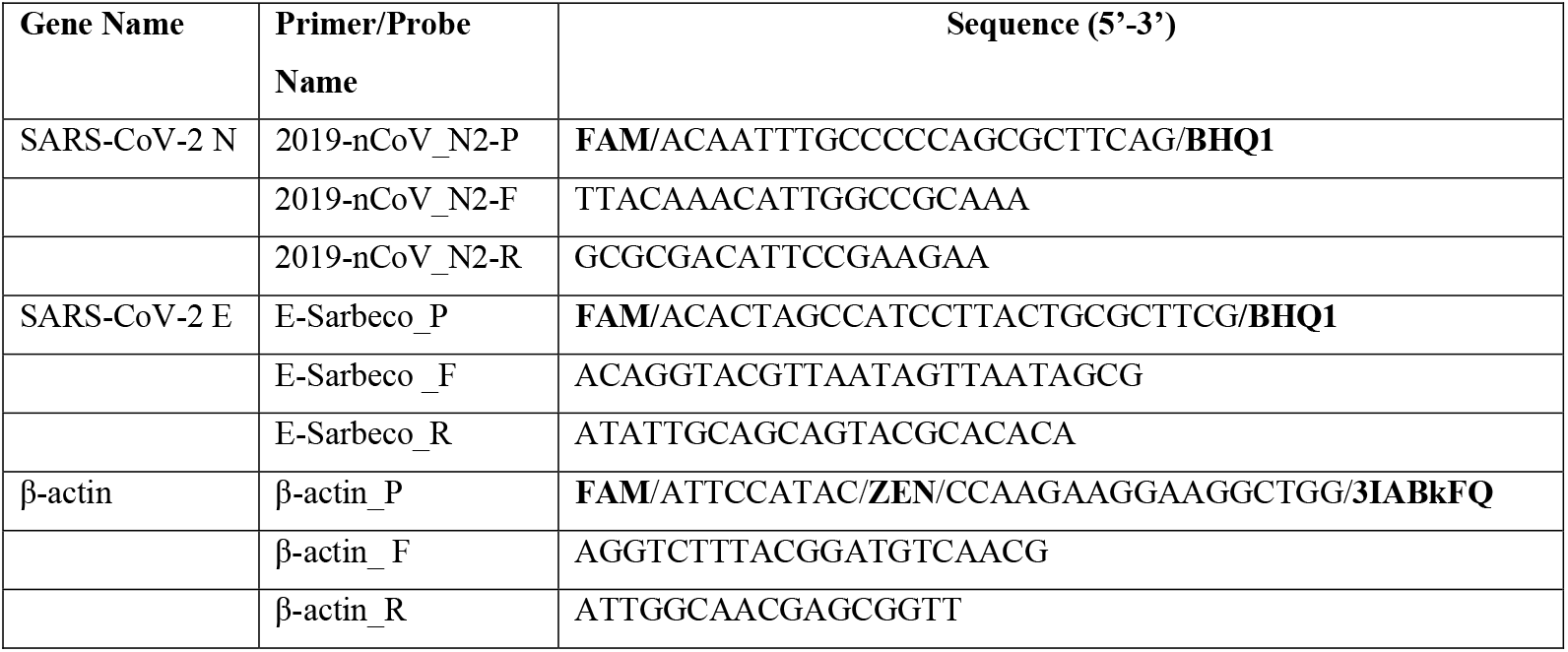
Probes and primers used in RT-qPCR assays.

Samples were run on an Applied Biosystems 7500 Fast PCR system (Thermo Fisher). The following cycling conditions were applied: reverse transcription for 10 min at 55 °C followed by denaturation at 95° C for 3 min. Then, 50 cycles of denaturation at 95 °C for 15 s and annealing/extension at 58 °C for 30 s. Amplification data were analized using Applied Biosystems 7500 software v2.3 (ThermoFisher). Results are reported as numbers of RNA copies for μg of total RNA.

### Statistical Analysis

When appropriate, data are presented as mean+standard deviation (SD). For the *in vivo* virus titration, virus dilutions and number of deaths/group after challenge were used to calculate the LD_50_ (Quest Graph™ LD_50_ Calculator, AAT Bioquest, Inc., S. Francisco, CA, USA). When indicated, the one- or two-tailed Mann–Whitney U test, the two-tailed Student T Test with Welch Correction, or the Kruskal-Wallis Test followed by Dunn’s Multiple Comparisons Test were conducted. Correlation analyses were conducted using the Spearman Rank Correlation Test. *p* <0.05 was considered statistically significant.

## Results

### Induction of N-specific CD8^+^ T polyfunctional and CD8^+^ T resident memory lymphocytes in lungs

K18-hACE2 transgenic mice were inoculated twice 2 weeks apart with vectors expressing either Nef^mut^ (6 mice) or Nef^mut^/SARS-CoV-2 N fusion protein (9 mice) following the experimental flow depicted in Fig. 1. From 10 to 15 days thereafter, N-specific CD8^+^ T cell immunity was evaluated in both circulating cells and lungs, in the latter case after intravenous labeling of a fluorescent anti-CD45 antibody to identify blood cells residing in both alveolar and interstitial compartments. IFN-γ EliSpot analysis for the detection of N-specific CD8^+^ T lymphocytes within circulating cells revealed a mean of more than 250 spots/2.5×10^5^ total splenocytes from vaccinated mice (Fig. 2A, Supplementary fig. 3), whereas mice injected with Nef^mut^-expressing DNA scored negative. By ICS/flow cytometry analysis, we found that immunization gave rise to a mean of 5.7% of N-specific CD8^+^ T lymphocytes, in the presence of a mean of more than 2% of polyfunctional (i.e., co-expressing IFN-γ, IL-2 and TNF-α) N-specific CD8^+^ T lymphocytes (Fig. 2B-D). When the ICS analysis was carried out in the CD45 negative fraction of cells isolated from lungs (identifying tissue-specific immune cells), higher percentages of both IFN-γ expressing and polyfunctional N-specific CD8^+^ T lymphocytes were found (Fig. 3A-C). Of importance, a well distinguishable population of N-specific Trm (i.e., CD44^+^, CD49a^+^, CD69^+^, CD103^+^, IFN-γ^+^) CD8^+^ T lymphocytes was found in lungs of vaccinated mice (Fig. 3D).

**Figure 1.**
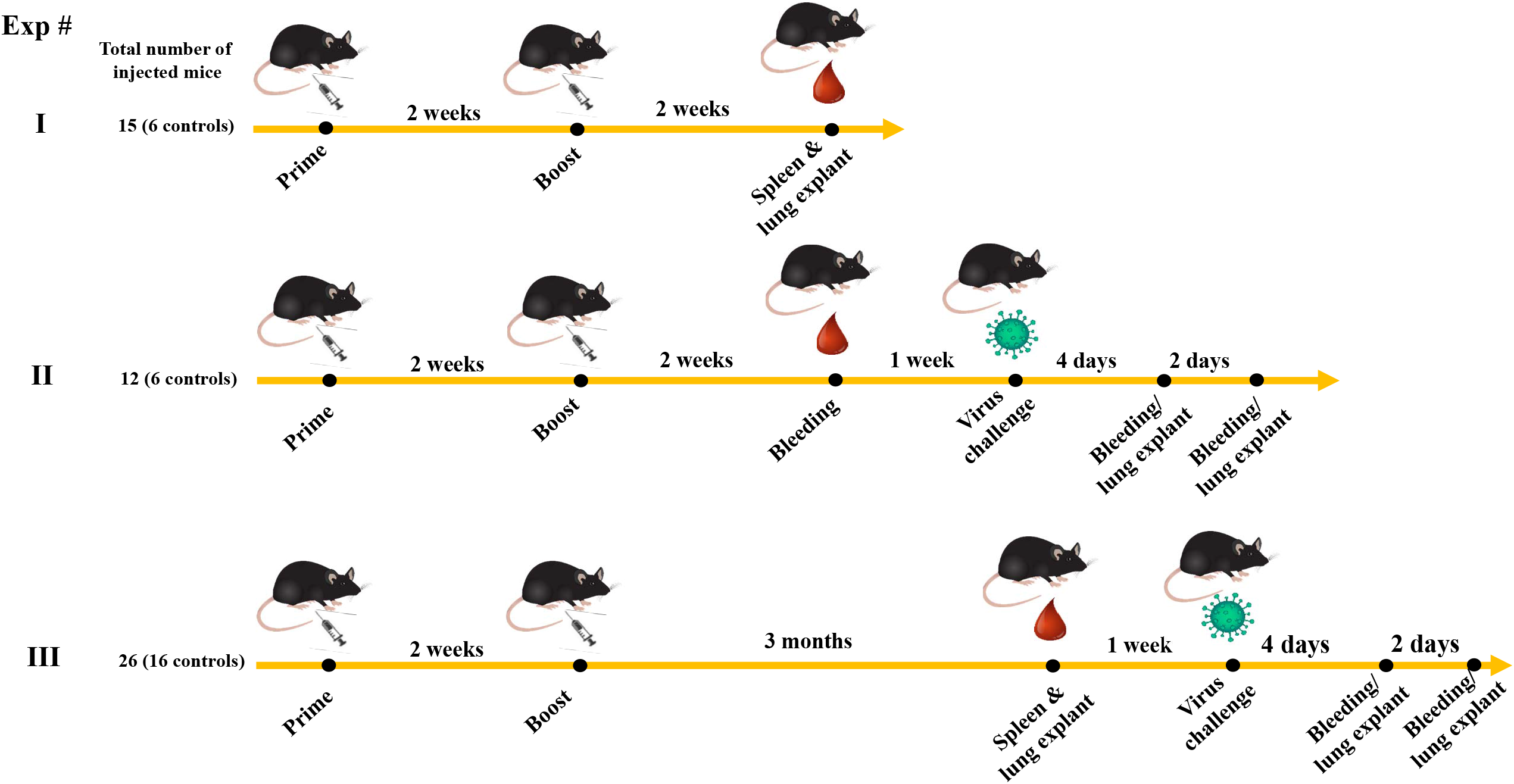
Scheme depicting both flow and time points of the here presented immunization/infection experiments carried out in K18-huACE2 mice. Shown is also the number of injected animal for each experiment.

**Figure 2.**
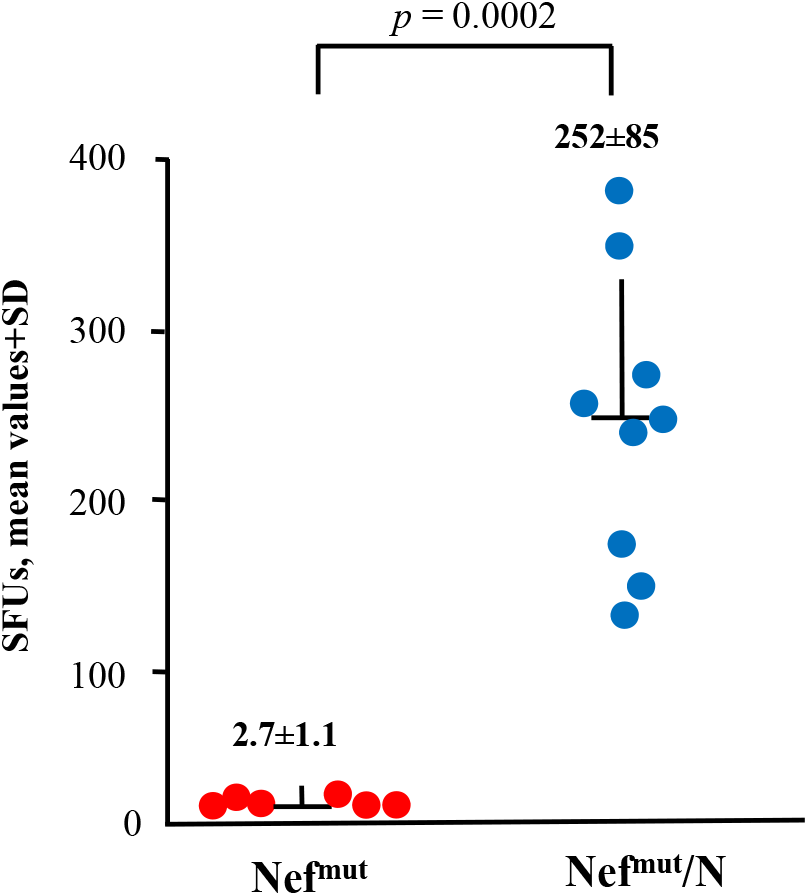
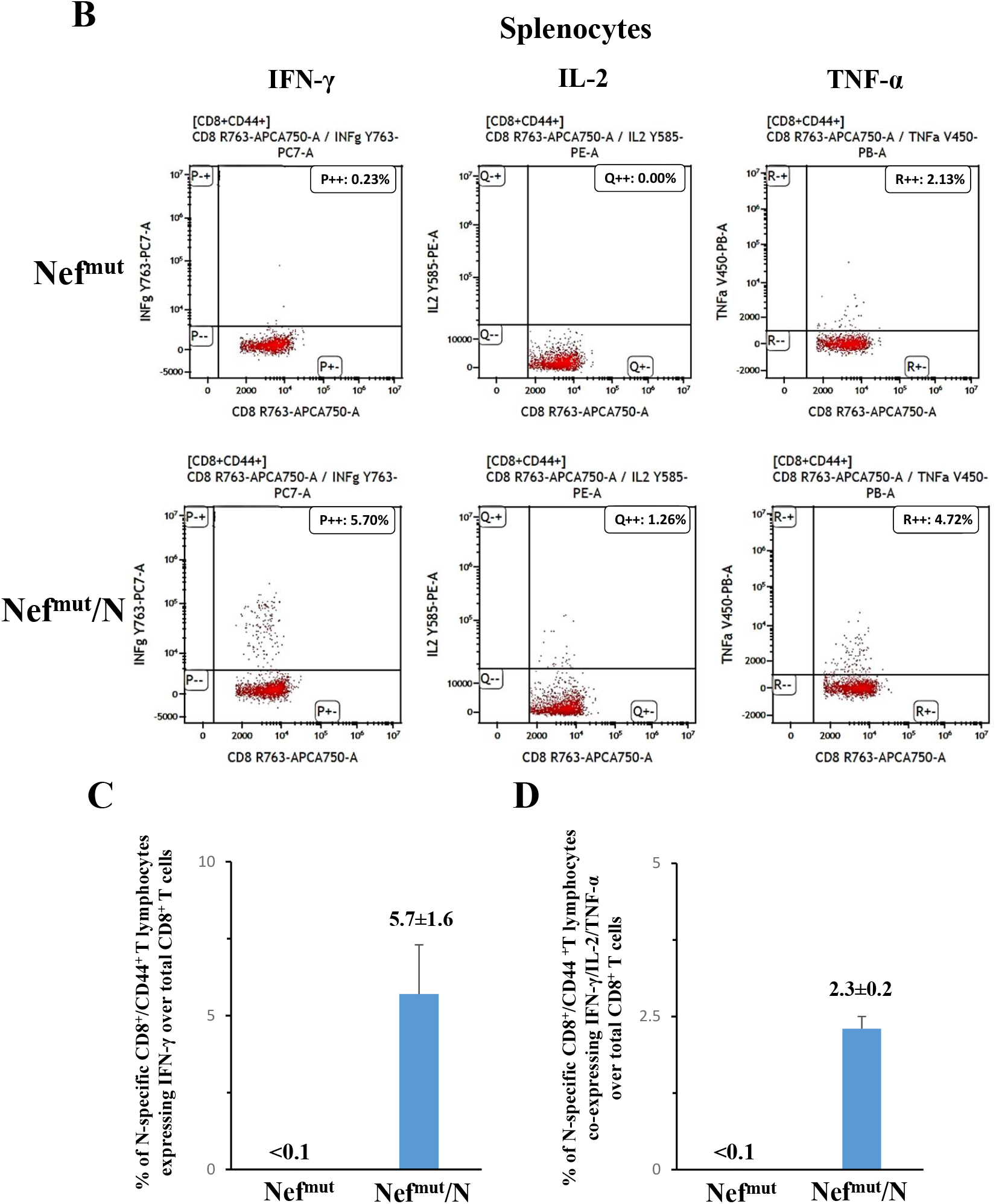
Detection of SARS-CoV-2-N-specific CD8^+^ T cells in splenocytes isolated from K18-hACE2 mice i.m. injected with either Nef^mut^- (6 mice) or Nef^mut^/N- (9 mice) expressing DNA vectors. **(A)** A total of 2.5×10^5^ splenocytes were incubated overnight with 5 μg/ml of either unrelated, Nef, or SARS-CoV-2-N specific peptides in IFN-γ EliSpot microwells. Shown are the numbers of spot-forming units (SFUs)/well calculated as mean values of triplicates after subtraction of mean spot numbers detected in wells of splenocytes treated with an unspecific peptide. Reported are intragroup mean values ±SD. Two-tailed Mann-Whitney U Test, *p* = 0.0002 **(B-D)** ICS/flow cytometry analysis of splenocytes isolated from mice injected with vectors expressing either Nef^mut^ or Nef^mut^/N. In panel **B**, raw data from a representative experiment for the detection of IFN-γ, IL-2, and TNF-α are presented. In panel **C**, shown are the percentages of cells expressing IFN-γ over the total of CD8^+^/CD44^+^ T cells from mice injected with the indicated DNA vectors. Mean values of the percentages of IFN-γ expressing cells from at least three pooled cell cultures treated with specific peptides after subtraction of values detected in cultures treated with an unrelated peptide are reported. In panel **D**, percentages of cells simultaneously expressing IFN-γ, IL-2, and TNF-α over the total of CD8^+^/CD44^+^ T cells are presented. Shown are mean values of the absolute percentages of cytokine expressing cells from at least three pooled cell cultures treated with specific peptides after subtraction of values detected in cultures treated with an unrelated peptide.

**Figure 3.**
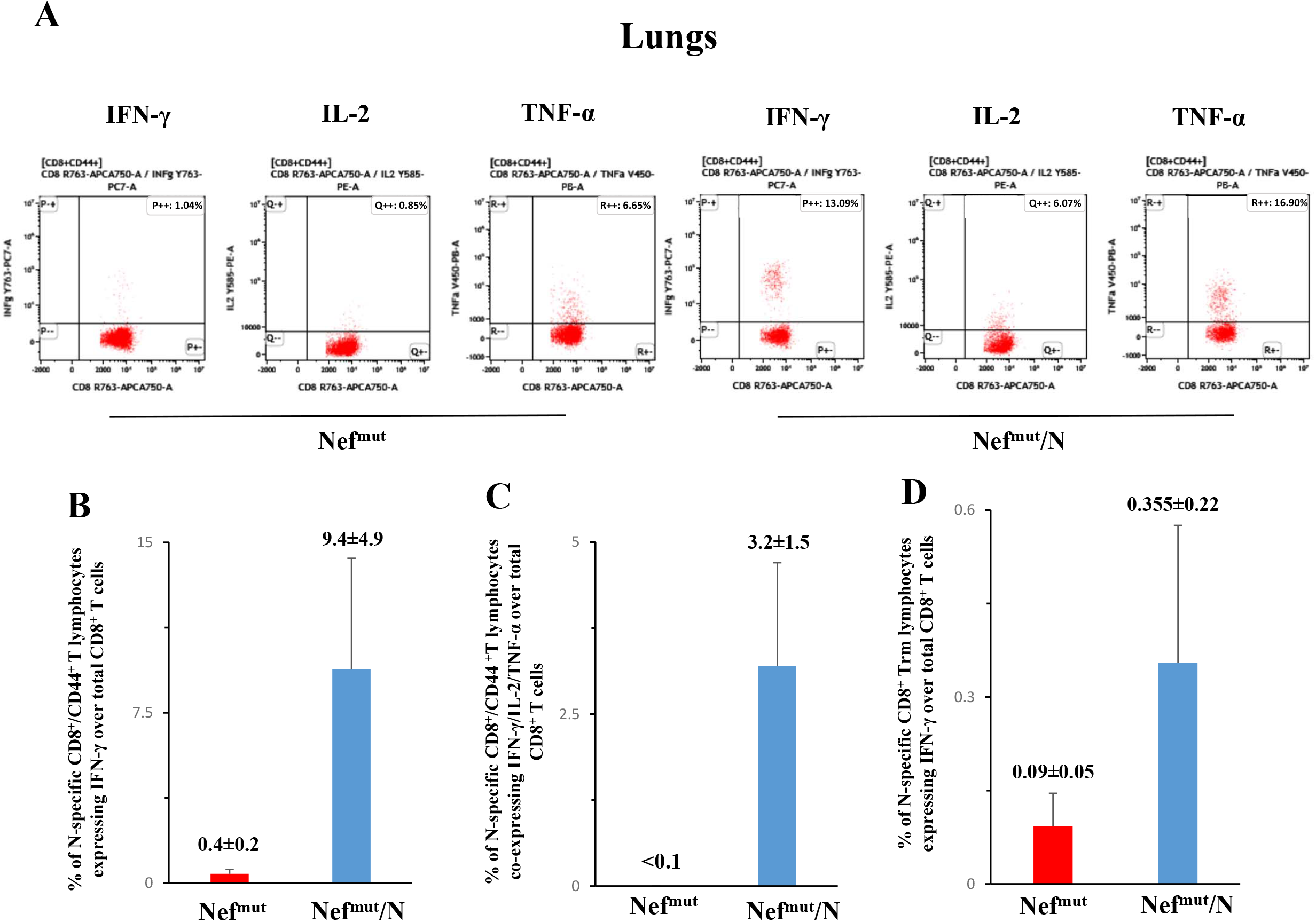
Detection of SARS-CoV-2-N-specific CD8^+^ T lymphocytes within immune cells isolated from lungs of K18-hACE2 mice i.m. injected with either Nef^mut^ or Nef^mut^/N-expressing DNA vectors. (**A**) Raw data from a representative ICS/flow cytometry analysis for the detection of IFN-γ, IL-2, and TNF-α within the CD45^-^ fraction of cells isolated from lungs. **(B)** Percentages of cells expressing IFN-γ over the total of CD8^+^/CD44^+^ T cells within lung cells pooled from three mice injected with the indicated DNA vectors. Mean values ±SD of percentages of IFN-γ expressing cells from at least three pooled cell cultures treated with specific peptides after subtraction of values detected in cells treated with an unrelated peptide are presented. **(C)** Percentages of cells simultaneously expressing IFN-γ, IL-2, and TNF-α over the total of CD8^+^/CD44^+^ T cells within lung immune cells pooled from three mice injected with the indicated DNA vectors. Shown are mean values of the percentages ±SD of cytokine expressing cells from pooled cell cultures treated with specific peptides after subtraction of values detected in cultures treated with an unrelated peptide. **(D)** Percentages of CD8^+^ Trm cells expressing IFN-γ over the total of CD8^+^/CD44^+^ T lymphocytes. Shown are mean value of percentages ±SD of positive cells from pooled cell cultures treated with specific peptides after subtraction of values detected in cultures treated with an unrelated peptide.

We concluded that the immunization with Nef^mut^/N expressing vector induced a strong N-specific CD8^+^ T cell immunity in lung tissues associating with a sub-population of N-specific CD8^+^ Trm lymphocytes.

### Impaired SARS-CoV-2 replication in lungs of N-immunized mice

Actual effectiveness of any antiviral vaccine strategy should be evaluated in terms of its efficacy in reducing viral replication in the target tissue. In the case of SARS-CoV-2 infection, the assessment should be focused on lungs, considering that in humans Covid-19-related severe disease and death are correlated with an uncontrolled viral replication in pulmonary alveoli. To this aim, in a second set of immunization experiments mice (6 per group) were injected with either Nef^mut^ or Nef^mut^/N DNA vectors, and actual immunization was checked fifteen days after the second immunization by IFN-γ EliSpot assay on PBMCs (Fig. 4A). After additional seven days, mice were infected i.n. with 4.4 lethal doses of SARS-CoV-2, and lungs were isolated 4-6 days thereafter. Lung tissues were then processed for the extraction and purification of total RNA. RT-qPCR analysis was finally carried out to enumerate the copy numbers of both SARS-CoV-2 E- an N-related RNA molecules. Overall, we found that virus replicated in lungs of N-vaccinated mice from 100- to 150-fold less efficiently than in control mice (Fig. 4B; two-tailed Mann-Whitney U Test, *p* = 0.0022). Consistently with what previously reported in survival assays [14], the inhibition of virus replication appeared most effective (i.e., up to 3.5 logs of reduction) in mice that had developed stronger N-specific immune responses. In line with this observation, Spearman Rank Correlation Test showed a negative correlation between days 4 and 6-cumulative results of N-specific IFN-γ EliSpot and RT-qPCR assays on both E and N viral genes (Spearman R = -0.94, *p* = 0.0167, and Spearman R = -1.0, *p* = 0.0028, respectively).

**Figure 4.**
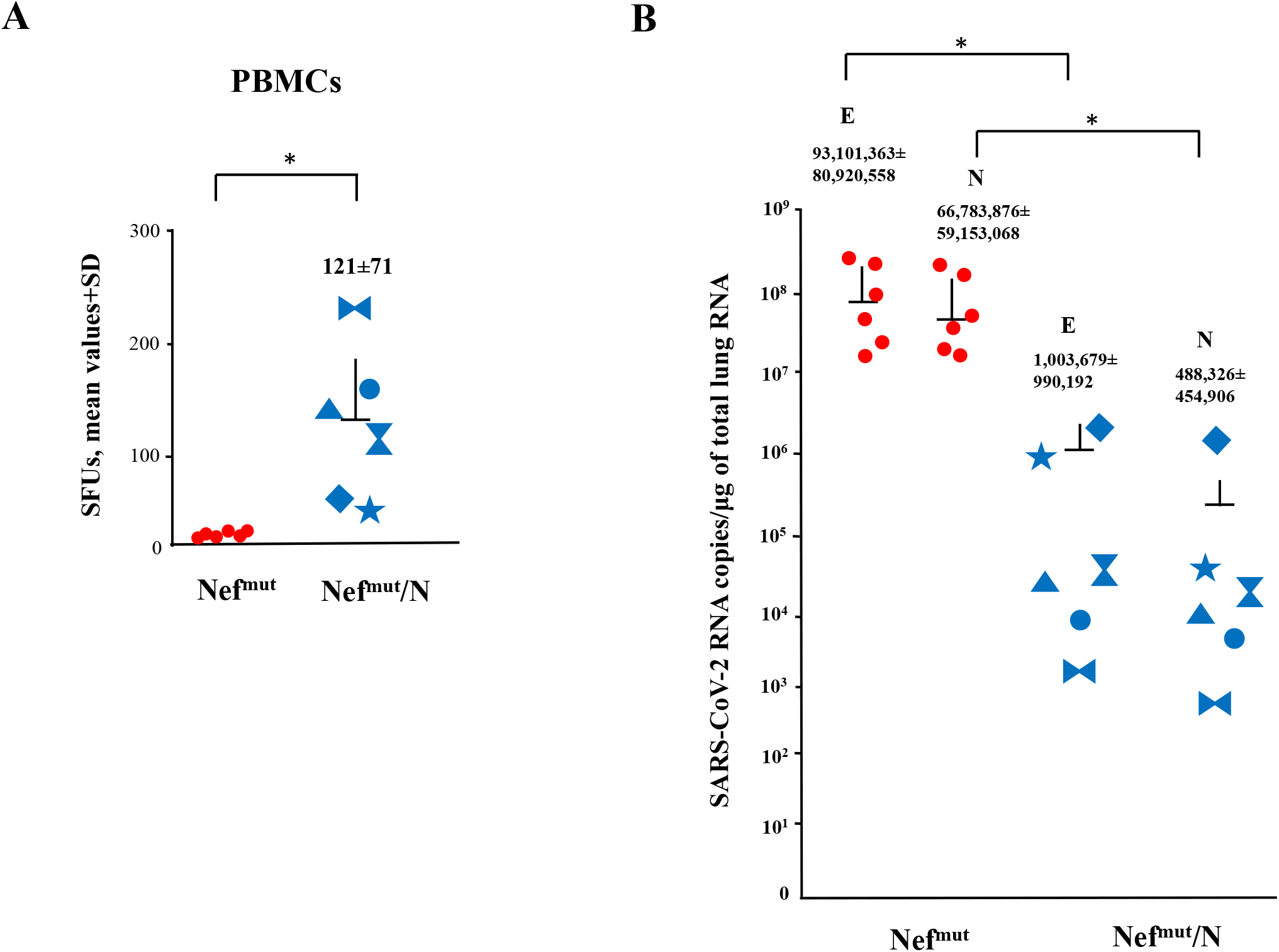
Viral loads in lungs of immunized mice. **(A)** Detection of N-specific immunity in vaccinated mice (6 mice per group). A total of 10^5^ PBMCs were incubated overnight with or without 5 μg/ml of either unrelated or SARS-CoV-2-N specific peptides in IFN-γ EliSpot microwells. Shown are the numbers of SFUs/well calculated as mean values of triplicates after subtraction of mean spot numbers detected in wells of splenocytes treated with an unspecific peptide. Reported is intragroup mean value ±SD. \**p* <0.05. **(B)** Viral loads in lungs of immunized/infected mice. Four and six days after challenges, injected mice (6 per group) were sacrificed, and lungs processed for the extraction of total RNA. One μg of total RNA from each infected mouse was then analyzed by RT-qPCR for the presence of both SARS-CoV-2 E- and N-specific RNAs. As internal control, actin RNA was also amplified. Shown are the numbers of viral RNA copies amplified from total RNA isolated from lungs of each animal, together with intragroup mean values ±SD. \**p* < 0.05.

### Intranasal SARS-CoV-2 infection does not affect the pre-existing N-specific cell immunity in lungs

It is well known that SARS-CoV-2 replication in lungs couples with hyper-inflammation and profound immune dysregulation [25-26]. Hence, we next were interested in monitoring whether and to what extent SARS-CoV-2 infection influenced the state of SARS-CoV-2-N-specific immunity in both circulating and lungs of vaccinated mice. To this aim, PBMCs recovered from infected mice were analyzed at both 4 and 6 days after challenge by both IFN-γ EliSpot and ICS/flow cytometry assays (Fig. 5A-B). We noticed that at day 6 post-infection, the Spike-specific CD8^+^ T cell immunity jumped dramatically in control mice, whereas it increased at apparently lower extents in N-vaccinated mice, in the presence of a persistent N-specific immunity. Analysis on cumulative day 4- and day 6-N-specific IFN-γ EliSpot assay results showed a statistically significant higher N-specific response in Nef^mut^/N vaccinated mice compared to control animals (Two-tailed Student T Test with Welch correction, *p* = 0.0015). When the ICS/flow cytometry analysis was carried out in cells isolated from lungs of infected mice at day 6 after challenge, in control mice striking levels of both Spike- and N-specific CD8^+^ T cell immunity were found, which conversely appeared moderate in N-immunized group. (Fig. 5C). Due to the limited number of recovered cells, cultures from lungs were set by pooling cells from three infected mice per group.

**Figure 5.**
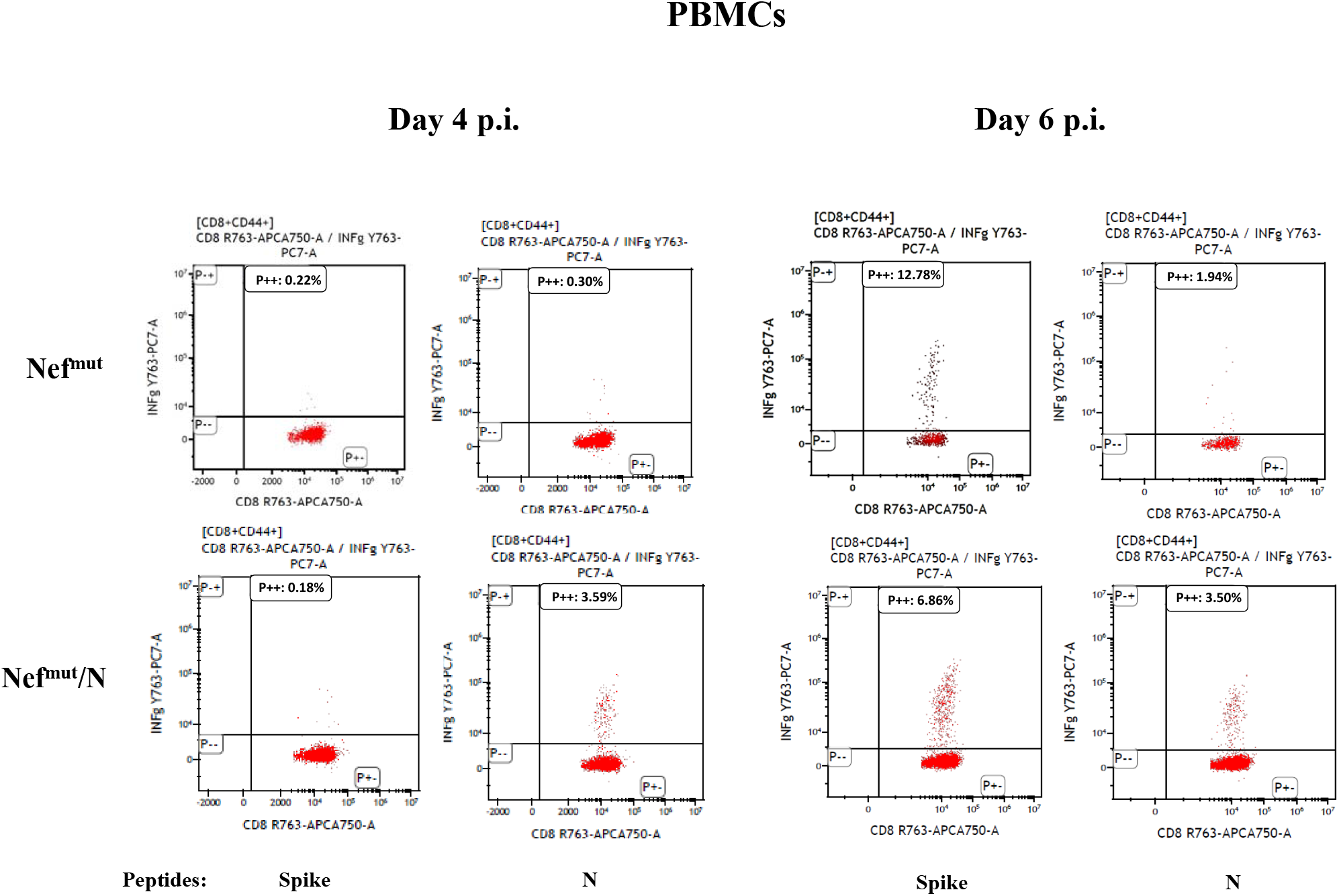
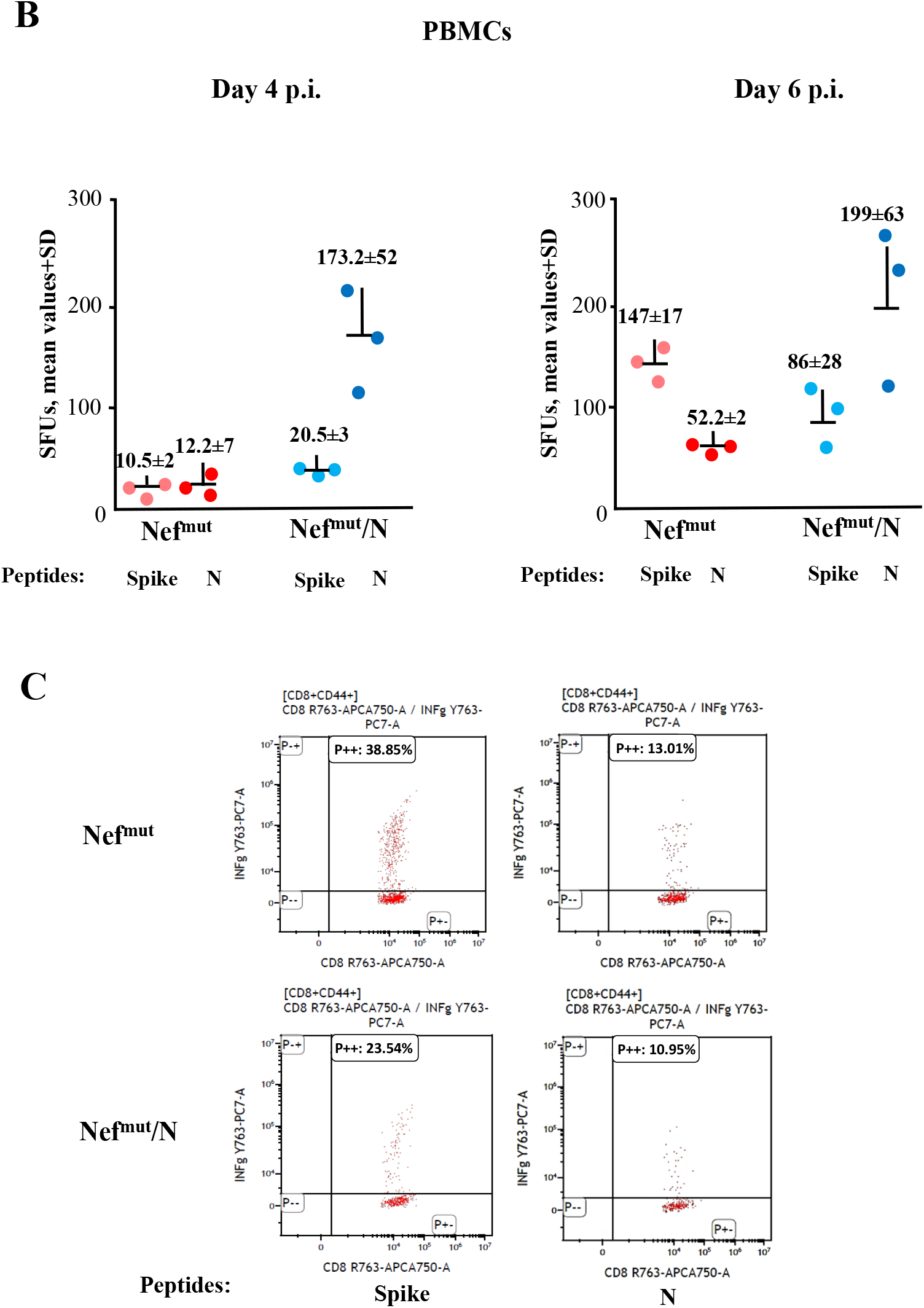
Detection of SARS-CoV-2-N-specific CD8^+^ T cells in K18-hACE2 mice i.m. injected with either Nef^mut^ or Nef^mut^/N-expressing DNA vectors and then infected. **(A)** Representative results from ICS/flow cytometry analysis for the expression of IFN-γ in PBMCs from K18-hACE2 mice injected with DNA expressing the indicated products, and then infected. Cells harvested 4 and 6 days after virus infection were cultivated overnight with either Spike-, N-specific, or unrelated peptides. Shown are data obtained with PBMCs pooled from three infected mice per condition representative of two experiments. Quadrants were set on the basis of cell fluorescence of samples treated with the unrelated peptide. **(B)** Detection by IFN-γ EliSpot analysis of both Spike- and N-specific CD8^+^ T cells within PBMCs from vaccinated mice both 4 and 6 days after infection. A total of 10^5^ PBMCs were incubated overnight with or without 5 μg/ml of either unrelated, Spike- or N-specific peptides in IFN-γ EliSpot microwells. Shown are the numbers of SFUs/well calculated as mean values of triplicates after subtraction of mean spot numbers calculated in wells of splenocytes treated with an unspecific peptide. Reported are intragroup mean values ±SD. **(C)** ICS/flow cytometry analysis for the detection of IFN-γ expressing CD8^+^ T lymphocytes within immune cells isolated 6 days after infection from lungs of K18-hACE2 mice i.m. injected with either Nef^mut^ or Nef^mut^/N-expressing DNA vectors. Cells pooled from lungs of three mice were incubated overnight with either Spike-, N-specific or unrelated peptides. Results are representative of two experiments. Quadrants were set on the basis of cell fluorescence of samples treated with the unrelated peptide.

These data indicated that the infection did not relevantly affect the levels of N-specific immunity in lungs of N-vaccinated mice.

### Persistent N-specific CD8^+^ T cell immunity in lungs of vaccinated mice

The persistence of immune protection over time is a mandatory feature for any vaccine strategy. In the case of EV-induced N-specific CD8^+^ T cell immunity, the prompt generation of an antigen-specific CD8^+^ Trm subpopulation in lungs was suggestive of a durable immune response. To test this hypothesis, the N-specific immune response was monitored three months after the second shot in a third set of immunization experiments. IFN-γ EliSpot analysis on splenocytes isolated from four mice per group showed the persistence of N-specific circulatory CD8^+^ T lymphocytes (Fig. 6A), although at levels reduced compared to what observed three weeks after the second injection. Most important, in these mice N-specific lung Trm scored more than 8% of total lung CD8^+^ Trm (Fig. 6B), with percentages over the total of CD8^+^ T lymphocytes apparently increased compared to those observed 2 weeks after boosters (Fig. 6C). These results were in line with the hypothesis that the injection of Nef^mut^/N DNA vector generated a long lasting immunity in lungs.

**Figure 6.**
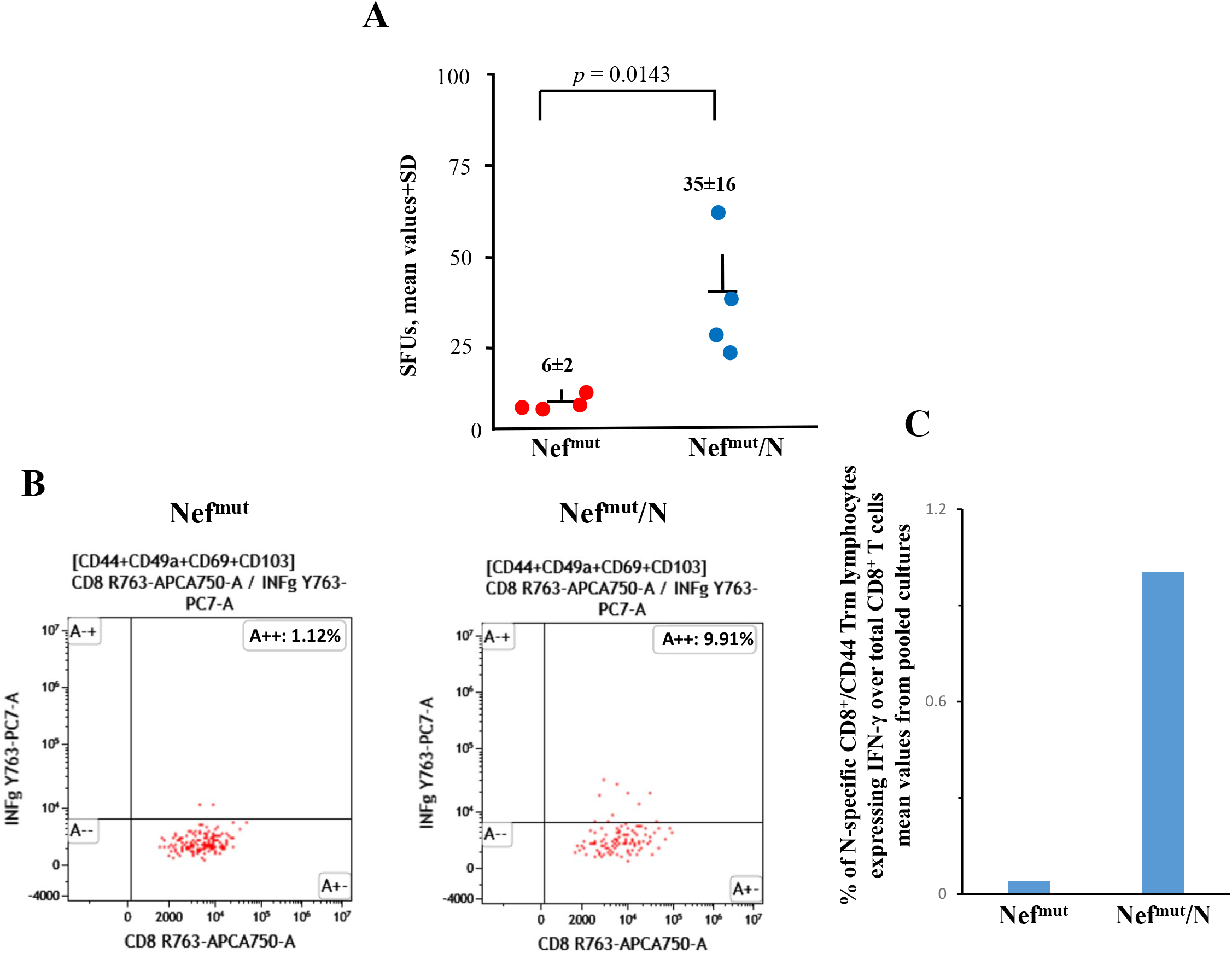
Detection of SARS-CoV-2-N-specific CD8^+^ T lymphocytes in K18-hACE2 mice three months after i.m. injection with either Nef^mut^ or Nef^mut^/N-expressing DNA vectors. **(A)** Detection of N-specific CD8^+^ T cells in injected mice. A total of 2.5×10^5^ splenocytes were incubated overnight with or without 5 μg/ml of either unrelated or SARS-CoV-2-N specific peptides in IFN-γ EliSpot microwells. Shown are the numbers of SFU/well calculated as mean values of triplicates after subtraction of mean spot numbers calculated in wells of splenocytes treated with an unspecific peptide. Reported are intragroup mean value ±SD. One-tailed Mann-Whitney U Test, *p* = 0.0143. **(B)** Raw data from a representative of two ICS/flow cytometry analysis for the detection of IFN-γ expressing CD8^+^ Trm lymphocytes within immune cells isolated from lungs of K18-hACE2 mice injected with either Nef^mut^ or Nef^mut^/N-expressing DNA vectors. **(C)** Percentages of CD8^+^ Trm cells expressing IFN-γ over the total of CD8^+^/CD44^+^ T lymphocytes isolated from lungs of mice injected with the indicated DNA vectors. Shown are mean values of percentages of positive cells from cultures treated with specific peptides after subtraction of values detected in cells treated with an unrelated peptide. The results are from two cell cultures each formed by the pool of cells from two immunized mice.

### Antiviral activity in lungs three months after immunization

Finally, we were interested to establish whether the persistence of N-specific Trm in lungs of vaccinated mice associated with an antiviral state. Six mice of the last set of immunization experiments were infected i.n. with identical experimental conditions used in previous challenges. In view of the previously described activation effect of Nef^mut^ on immature dendritic cells [27], which would favor the natural antiviral immune response, a group of six mice injected with saline solution (sham) was included in the challenge experiment. The RT-qPCR analysis of total RNA isolated from lungs 4-6 days after challenge showed an overall drop of virus replication extents in N-immunized mice compared to controls (Fig. 7A) similar to that observed three weeks after boost. Kruskal-Wallis Test followed by Dunn’s Multiple Comparisons Test showed a statistically significant control of infection in Nef^mut^/N-vaccinated mice compared to the sham group in terms of both measured E and N viral genes, with *p* < 0.01 for both. Interestingly, the immune response induced by Nef^mut^ seemed to contribute to the overall antiviral effect induced by the immunization against Nef^mut^/N. As assessed by IFN-γ EliSpot analysis carried out on PBMCs isolated at both 4 and 6 days after challenge, the Spike-related CD8^+^ T cell immune responses were potently generated after infection of control mice. On the other hand, the Spike-specific CD8^+^ T cell immune response in N-immunized mice was elicited at lower levels compared to control mice (Kruskal-Wallis Test followed by Dunn’s Multiple Comparisons Test, Nef^mut^/N-vaccinees vs sham mice, at day 6: *p* < 0.05), in the presence of an N-specific immunity not diminished compared to pre-infection levels (Fig. 7B).

**Figure 7.**
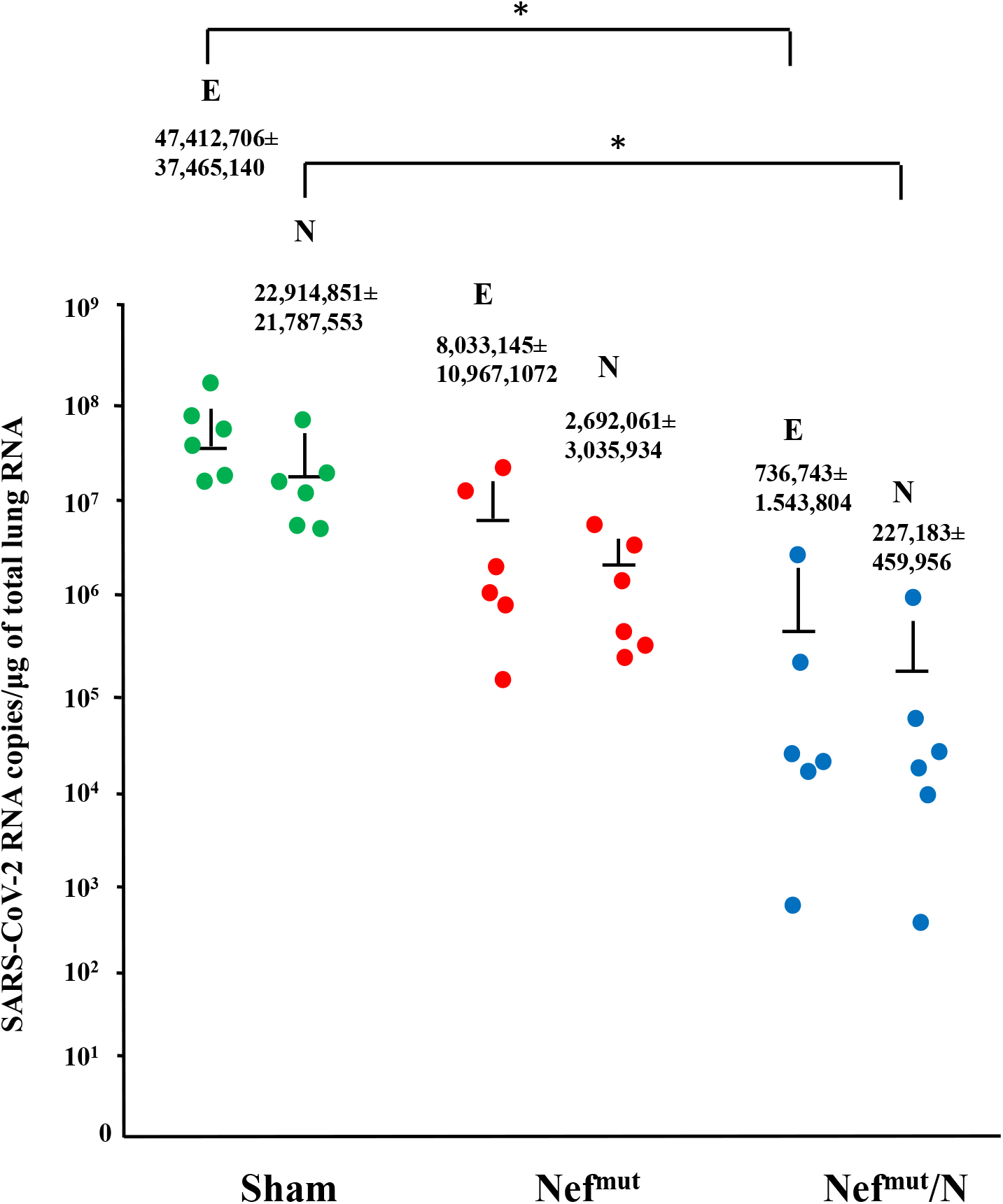
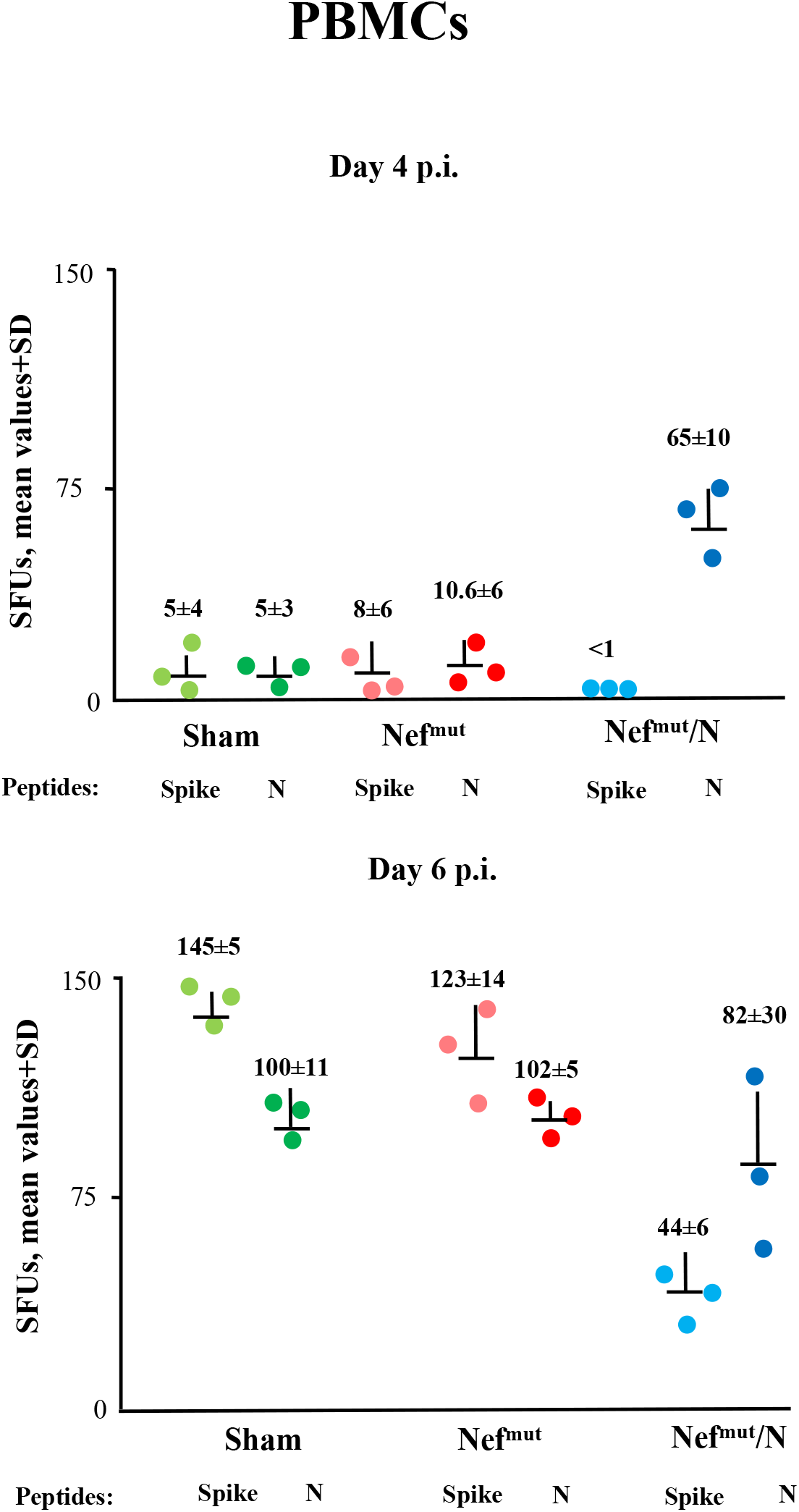
Viral loads in lungs of mice infected three months after last injections. **(A)** RT-qPCR analysis. Four to **s**ix days after challenge, injected mice (6 per group) were sacrificed, and lungs processed for the extraction of total RNA. One μg of total RNA from each infected mouse was then analyzed by RT-qPCR for the presence of both SARS-CoV-2 E- and N-specific RNAs. As internal control, actin RNA was also amplified. Shown are the numbers of viral RNA copies amplified from total RNA isolated from lungs of each animal, together with intragroup mean values ±SD. \**p* < 0.01. **(B)** Detection of both Spike- and N-specific CD8^+^ T cells within PBMCs isolated both 4 and 6 days after infection. A total of 10^5^ PBMCs were incubated overnight with or without 5 μg/ml of either unrelated, Spike- or N-specific peptides in IFN-γ EliSpot microwells. Shown are the numbers of SFUs/well calculated as mean values of triplicates after subtraction of mean spot numbers calculated in wells of splenocytes treated with unspecific peptides. Reported are intragroup mean values ±SD.

Taken together, these data support the idea that CD8^+^ T N-specific immunity induced in lungs is able to control virus replication in a durable manner.

## Discussion

Respiratory airways are the battlefield where the fate of subjects infected by SARS-CoV-2 is determined. The efficacy of anti-SARS-CoV-2 vaccines would be the result of promptness, potency, and durability of the immune response evoked in both upper and lower respiratory tracts. In humans, current mRNA-based vaccine have been shown to induce immune responses quantitatively limited and qualitatively incomplete in both tracts [28]. Second-generation vaccines are expected to overcome such limitations by focusing their immunogenic activity at pulmonary level. In this perspective, we investigated the potentialities of an innovative anti-SARS-CoV-2 CD8^+^ T cell-based vaccine consisting in engineered EVs spontaneously released by muscle cells incorporating the SARS-CoV-2 N protein. They are produced upon i.m. injection of a DNA vector expressing the SARS-CoV-2 N protein fused at the C-terminus of Nef^mut^, i.e., an EV-anchoring protein incorporating in EVs at very high efficiency even when fused with heterologous proteins [29]. Of importance, we previously demonstrated that the Nef^mut^-dependent antigen incorporation into EVs is critical for the induction of the strong CD8^+^ T cell immunity, which does not take place when the antigen is expressed alone [30].

This model has been investigated in survival assays on K18-hACE2 transgenic mice using SARS-CoV-2 N engineered EVs as immunogen [14]. However, such an animal model recapitulates the Covid-19 pathogenesis in humans only partly. In fact, results from different papers demonstrated that, upon i.n. challenge, the most evident clinical signs of infection derive almost exclusively from the viral neuroinvasion, with cells of olfactory bulbs acting as viral port of entry [16-19]. Hence, evaluating the efficacy of anti-SARS-CoV-2 compounds in K18-hACE2 transgenic mice by measuring weight loss, ataxia, and death could lead to results of not obvious significance for humans. Conversely, monitoring the antiviral effects in terms of virus replication extents in lungs would be much more informative for a possible translation in humans, since SARS-CoV-2 replicates efficiently in lungs of humans as well [16].

Using fluorescent EVs, several authors demonstrated that circulating EVs diffuse in lungs quite efficiently [31-34]. We assume that engineered EVs emerging from muscle cells expressing Nef^mut^/N fusion products freely circulate into the body reaching mediastinal lymph nodes and lung germinal centers, where N-specific CD8^+^ T lymphocytes are selected and activated, with generation of N-specific CD8^+^ Trm. This mechanism might explain how a distal i.m. injection could result in a sustained cell immunity in lungs.

We measured the viral replication extent in lungs of vaccinated mice through RT-qPCR analysis. Several authors reported that the i.n. infection of K18-huACE2 transgenic mice induces a peak of virus replication in lungs at days 2-3 post infection. No major differences in the extents of viral replication were observed at days 4-7 post challenge [16, 18, 35, 36]. These well consolidated evidences justified the gathering of our data obtained from lungs excised at days 4 and 6 after infection. In evaluating these data, it should be considered that, despite its high sensitivity, the assay could not distinguish RNA viral molecules incorporated into infectious viral particles from those associated with non-infectious ones. On the other hand, the assay cannot discriminate at what extent cell-associated viral RNA molecules will be allocated in emerging viral particles. In addition, one should consider the antiviral mechanism on the basis of the CD8^+^ T cell immunity. Differently from neutralizing antibodies, which prevent virus cell entry, virus-specific CD8^+^ T lymphocytes can control viral replication by recognizing and destroying cells exposing virus-related peptides on MHC Class I, thus when high levels of intracellular viral RNA have been already produced.

RT-qPCR data we obtained in mice challenged 3 months after immunization suggested that the response to Nef^mut^ alone might be part of the overall antiviral effect observed in mice injected with Nef^mut^/N. In view of the previously documented activation effect of Nef^mut^ on dendritic cells [27], it is tempting to speculate that the overall N-specific CD8^+^ T cell immune responses, i.e., natural and vaccine-induced, would be favored by a sort of persistent pre-activation state of pulmonary dendritic cells induced by Nef^mut^ upon internalization of engineered EVs. Notably, in light of the results obtained three months after boost, it cannot be excluded that the Nef^mut^/N-induced antiviral effect observed 3 weeks after last immunizations was underestimated.

We noticed that the levels of emerging Spike-specific CD8^+^ T lymphocytes in lungs appeared reduced in vaccinated mice compared to control ones. We assumed that this effect was consequence of the inhibitory effect on viral replication. On the other hand, the detection of a quantitatively similar N-specific sub-population of CD8^+^ Trm lymphocytes 3 weeks and 3 months after boosting was strongly suggestive of the establishment of a long lasting anti-SARS-CoV-2 immunity at lungs. The strong reduction of viral replication extents we detected 3 months after vaccination was in line with this idea. Considering that it was largely demonstrated that lung CD8^+^ Trm can duplicate even in absence of antigenic stimulus [37], it is tempting to speculate that this antiviral immunity has the potential to persist for a very long time. Based on these evidences, lung N-specific CD8^+^ Trm might be considered the most appropriate immunologic correlate of the observed inhibition of viral replication in lungs.

Our study present a number of limitations. Virus from lungs of challenged mice was not titrated in terms of infectious units. The antiviral effect in lungs of vaccinated mice was evaluated using a single, very high viral dose, i.e., 4.4. LD_50_. In addition, dose-response experiments with different amounts of injected DNA are lacked. Finally, we cannot exclude that an N-specific CD4^+^ T lymphocyte response in lungs can contribute to the overall antiviral effect.

Orally administrated vaccines would represent a relevant complementation/amelioration of current vaccines, which are not effective enough to impede virus replication in respiratory mucosa [38-42]. Here presented results, together with those very recently published regarding the immunogenicity of SARS-CoV-2 N-engineered EVs in *ex vivo* human cells [43], can be instrumental to design a second-generation, innovative anti-Covid-19 vaccine based on mucosal administration of N-engineered EVs. In this way, optimal levels of CD8^+^ T cell immunity should be achieved at the viral port of entry.

## Author Contributions

Conceptualization, M.F., F.M. C.C; methodology, M.A., M.S., A.D., A.G., P.L., K.P., Z.M.; investigation: F.M., C.C., F.F, M.A.; resources: M.V., A.C., M.F.; data curation: F.M., C.C., F.F., M.A.; writing-original draft preparation: M.F.; writing-review and editing: F.F., C.C., F.M.; funding acquisition: M.F.

## Institutional Review Board Statement

The study was conducted according to the guidelines of the Declaration of Helsinki, and approved by the Institutional Review Board of the Istituto Superiore di Sanità (protocol code n. 565/2020 PR, approved on June 6, 2020, and protocol code 591/2021 released on and July 30^th^ 2021).

## Informed Consent Statement

Not applicable

## Data Availability Statement

The data presented in this study are available on request from the corresponding author. The data are not publicly available due to patent application.

## Acknowledgments

Both SARS-Related Coronavirus 2, Isolate Italy-INMI1, NR-52284, and the peptide array of SARS-Related Coronavirus 2 Spike Glycoprotein, NR-52402, were obtained through BEI Resources, NIAID, NIH. We thank Antonio Di Virgilio, Patrizia Cocco, Ferdinando Costa, Daniela Diamanti, Fabiola Diamanti, and Pietro Arciero Istituto Superiore di Sanità, for technical support, and Federica Magnani and Rosangela Duranti Istituto Superiore di Sanità for secretarial assistance.

---

**Supplementary fig. 1.**
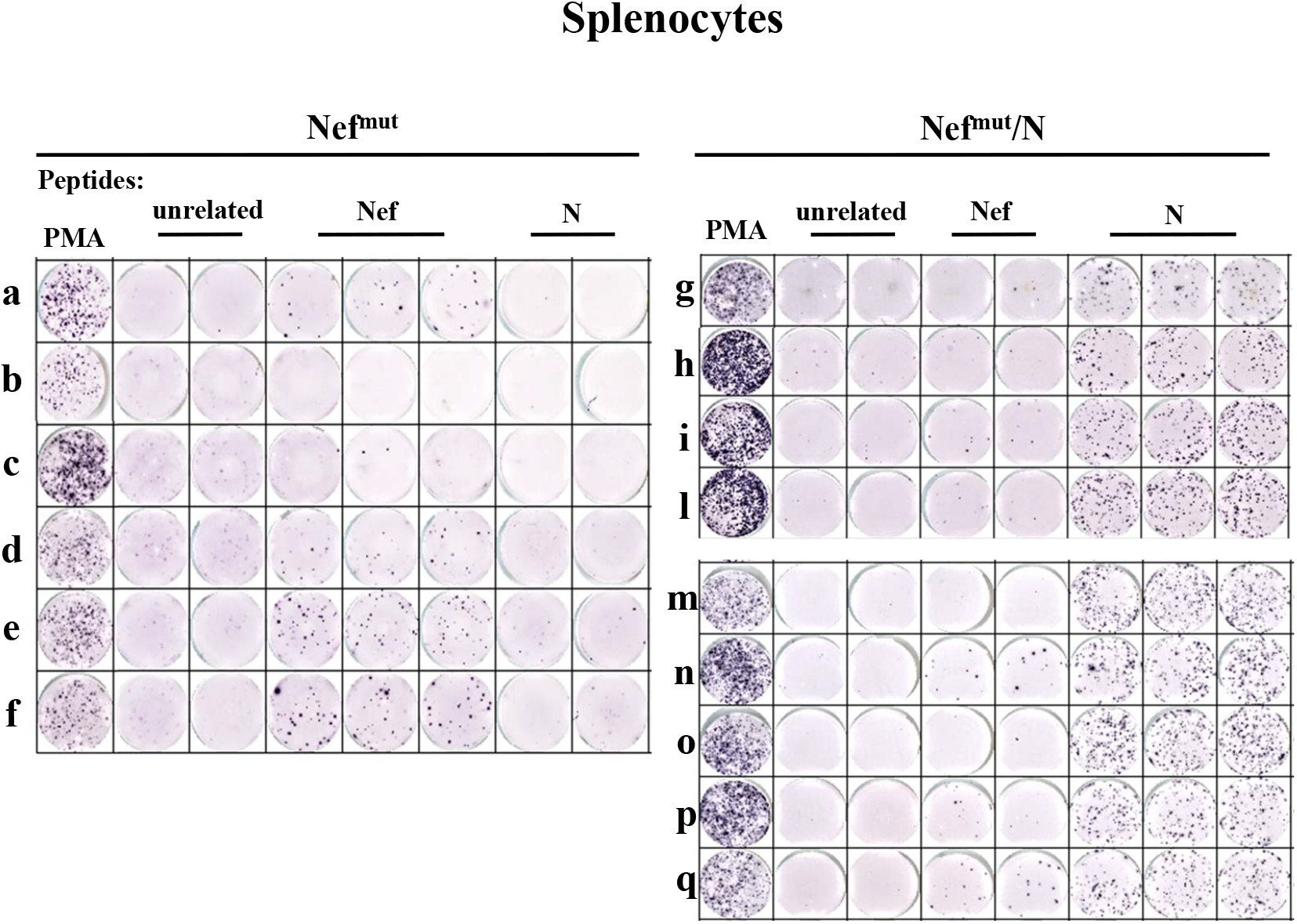
Raw data from IFN-γ EliSpot analysis for the detection of SARS-CoV-2-N-specific CD8^+^ T cells in splenocytes isolated from K18-hACE2 mice i.m. injected with either Nef_mut_- (6 mice, a-f) or Nef_mut_/N- (9 mice, g-q) expressing DNA vectors. A total of 2.5×10^5^ splenocytes were incubated overnight with 5 μg/ml of either unrelated, Nef, or SARS-CoV-2-N specific peptides in IFN-γ EliSpot microwells. As control, cells were also treated with PMA and ionomycin.

**Supplementary fig. 2.**
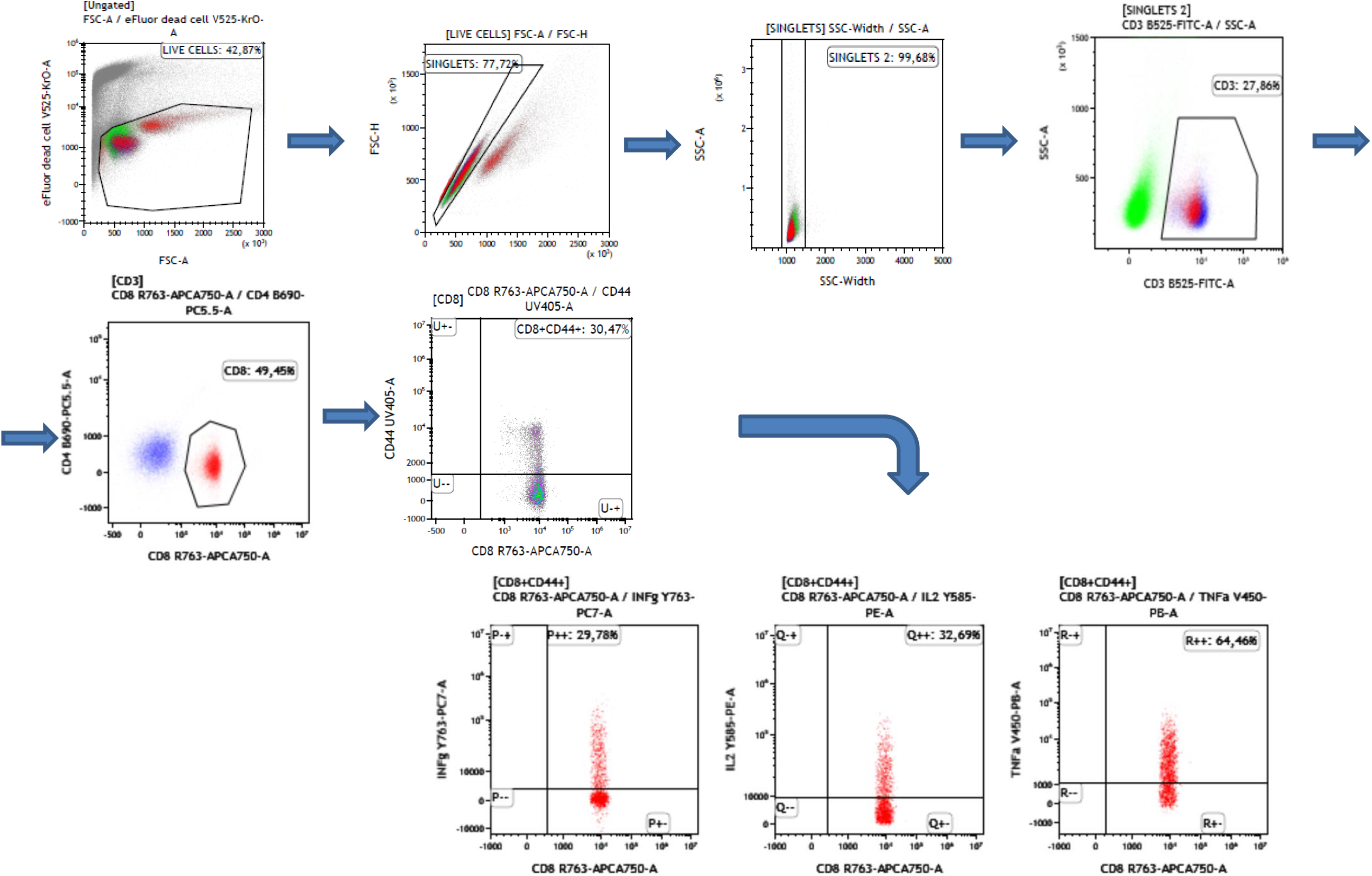
Gating strategy carried out in ICS/flow cytometry analysis of splenocytes from injected mice. Shown is the analysis of PMA-treated cells.

**Supplementary fig. 3.**
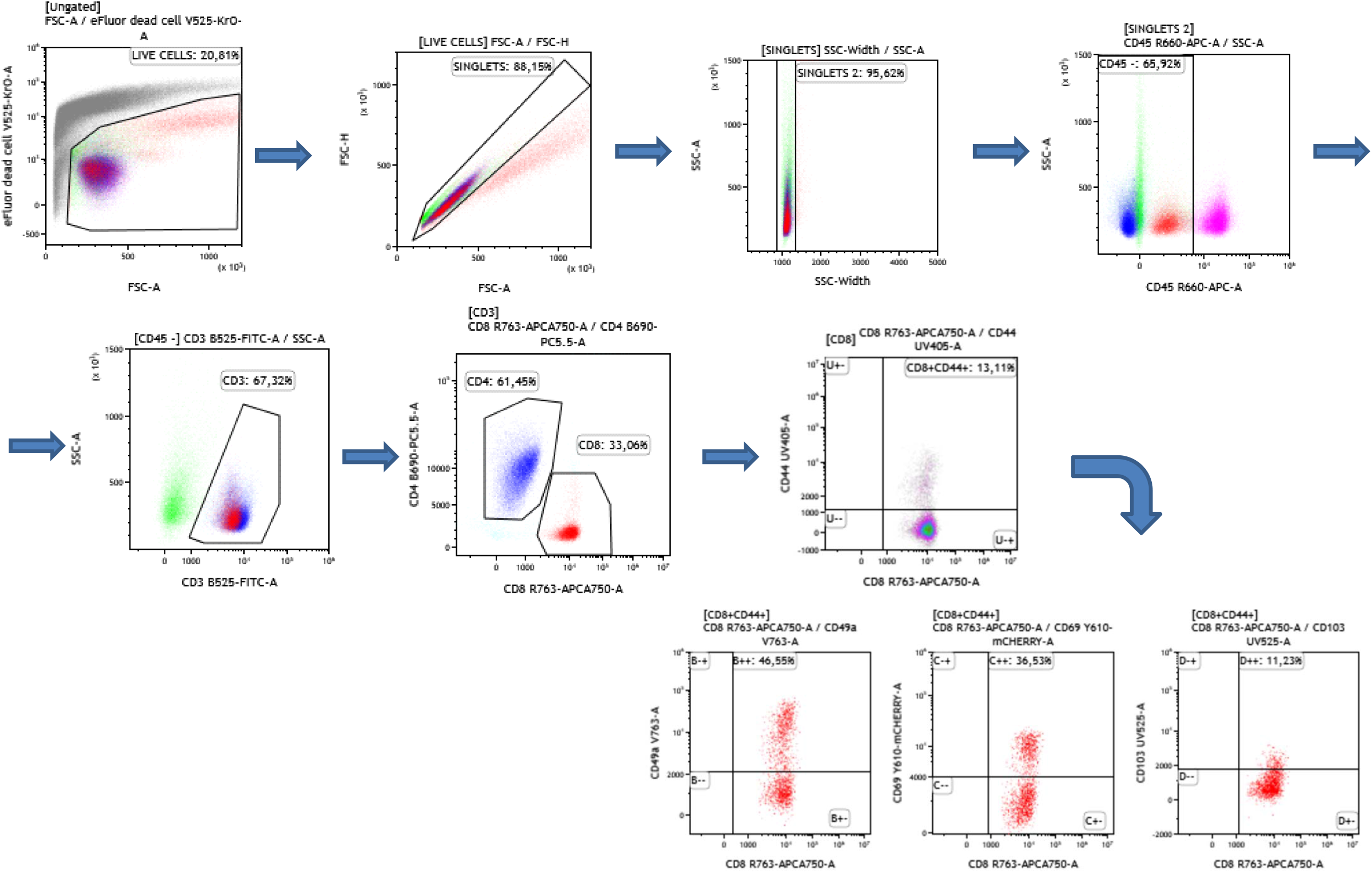
Gating strategy carried out in ICS/flow cytometry analysis of CD45 negative immune cells isolated from lungs of injected mice. Shown is the analysis on cells treated with an unrelated peptide

